# Co-localization and confinement of diphosphohydrolases and ecto-nucleotidases modulate extracellular adenosine nucleotide pools

**DOI:** 10.1101/670570

**Authors:** Rahmaninejad, Pace, Bhatt, Sun, Kekenes-Huskey

## Abstract

Nucleotides comprise small molecules that perform critical signaling and energetic roles in biological systems. Of these, the concentrations of adenosine and its derivatives, including adenosine tri-, di-, and mono-phosphate are dynamically controlled in the extracellular-space by diphosphohydrolases and ecto-nucleotidases that rapidly degrade such nucleotides. In many instances, the close coupling between cells such as those in synaptic junctions yields tiny extracellular ‘nanodomains’, within which the charged nucleotides interact with densely-packed membranes and biomolecules. While the contributions of electrostatic and steric interactions within such nanodomains are known to shape diffusion-limited reaction rates, less is understood about how these factors control the kinetics of sequentially-coupled diphosphohydrolase/nucleotidase-catalyzed reactions. To rank the relative importance of these factors, we utilize reaction-diffusion numerical simulations to systematically probe coupled enzyme activity in narrow junctions. We perform these simulations in nanoscale geometries representative of narrow extracellular compartments, within which we localize sequentially- and spatially-coupled enzymes. These enzymes catalyze the conversion of a representative charged substrate such as adenosine triphosphate (ATP) into substrates with different net charges, such as adenosine monophosphate (AMP) and adenosine (Ado). Our modeling approach considers electrostatic interactions of diffusing, charged substrates with extracellular membranes, and coupled enzymes. With this model, we find that 1) Reaction rates exhibited confinement effects, namely reduced reaction rates relative to bulk, that were most pronounced when the enzyme was close to the pore size and 2) The presence of charge on the pore boundary further tunes reaction rates by controlling the pooling of substrate near the reactive protein akin to ions near trans-membrane proteins. These findings suggest how remarkable reaction efficiencies of coupled enzymatic processes can be supported in charged and spatially-confined volumes of extracellular spaces.

## 2 Introduction

Nucleotide signaling and regulation of cellular energy pools are reliant on the diffusion of small molecules over micrometer-scale distances [1]. Examples of processes reliant on nucleotides include signal transduction and regulation in smooth muscle [2], network motifs in transcriptional regulation networks [3], genomic regulatory networks [4], complexes of metabolic enzymes [5] and trans-membrane ligand-gated channels [6, 7]. Of the latter, many nucleotide-gated channels and ATPases [8] reside within extracellular junctions formed between cells in close apposition [9], such as synaptic junctions comprised of neurons and glia [10]. Scanning electron microscopy has revealed that many of these junctions are on the nanometer length-scale [11]. Within those spaces, we expect that the free diffusion and apparent concentration of nucleotides will differ substantially from bulk solutions, although the influence of electrostatics, and confinement on diffusion properties and distributions of nucleotides have only been examined in limited detail.

At the cell surface, nucleotide pools are controlled by phosphohydrolases and nucleotidases [12, 13]. These enzymes hydrolyze nucleotides including adenosine- and uracil-based molecules and thereby regulate the pool of nucleotides available for signaling and metabolism [14]. It has for instance been demonstrated that the nucleotide concentration can vary considerably at the cell surface on both cytoplasmic and extracellular sides, as measured by the activity of ATPases and ATP-sensitive channels [15, 16]. A sub-class of phosphohydrolases called ecto-nucleotidase (NDA)s are localized to the extracellular surfaces of cell membranes. There, NDAs rapidly and dynamically control nucleotide concentrations adjacent to proteins that catalyze or are gated by these molecules. These include proteins such as purinergic receptors triggered by ATP and adenosine diphosphate (ADP)binding [14]. Although many NDAs are relatively nonspecific in their affinities for adenosine phosphates, some classes are selective for ATP and ADP, such as CD39a and CD39b, respectively [12, 17]. Subsequently, AMP hydrolysis into adenosine proceeds via the CD78 ecto-nucleotidase [12]. In this capacity, CD39 and CD78 catalyze the coupled, sequential hydrolysis of ATP into adenosine, and thereby influence nucleotide signaling at the membrane. However, no precise characterization is available for ATP diffusion-limited reaction kinetics in nucleotide pools localized to the membrane as a result of these enzymes’ spatial configurations within junctions. Thus, the pools that are actually encountered by proteins remain poorly resolved.

A key foothold for understanding ATP pools is to examine factors that control the coupling of sequential reactions, such as *ATP → AMP* and *AMP → Ado*. The fundamental motif of a sequentially-controlled enzymatic process consists of two enzymes, of which one enzyme generates a reaction intermediate that is catalyzed by the second enzyme [5, 18]. For diffusion-limited reactions, the efficiency of sequentially-coupled reactions is strongly determined by the relative distance between enzymes, as well as the rates of substrate diffusion toward their reactive centers [19]. Further, sequential enzyme reactivity depends on the transfer efficiency of intermediates, which can be facilitated by molecular tunnels [20] or electrostatic channeling [19, 21]. An essential consideration is therefore how intrinsic rates of substrate diffusion in bulk solution are modulated by steric and long-range electrostatic interactions between substrates and target enzymes versus those with the surrounding cellular environment [22, 23]. For instance, nucleotides are generally negatively-charged and are thus attracted to positively-charged nucleotide binding sites of NDA[24]. Most notably, cellular ‘crowders’ comprising enzymes, proteins, and macromolecules typically reduce substrates’ intrinsic diffusion rates, which in turn can manifest in altered enzyme kinetics [23, 25–29]. Additionally, diffusion limitations stemming from densely packed media or impermeable membranes can confine substrates to narrow ‘microdomains’, within which substrate concentrations are vastly different from those in the bulk cytosol or extracellular medium [30]. Based on these considerations, it is plausible that nucleotidases confined to narrow compartments between cells will hydrolyze extant nucleotide pools with markedly different kinetics than those observed *in vitro*. However, only recently has coupled NDA activity been examined [31] via numerical modeling; in that study, Sandefur *et al* [31] developed a computational model of human airway surface hydration that accurately described experimentally-observed trends in NDA activity. This study was an important first step toward describing coupled NDA enzyme activities. Extension of this description to account for the extracellular environment between cells, including diffusion-limitation imposed by macromolecules and long-range electrostatic interactions from charged proteins and membranes are expected to give local variations that strongly differ from bulk measurements.

Reaction kinetics in biological media are inherently difficult to study, given the breadth of influential factors including weak interactions of substrates with lipid membranes or proteins, restricted accessible volumes owing to such crowding by adjacent enzymes, proximity between enzymes involved in catalysis, and long-range electrostatic interactions. Since the systematic control of these factors in experiments is challenging [32], numerical models of molecular diffusion and reaction kinetics have been valuable in our understanding of catalysis in biological systems. At the coarsest resolution of such numerical models are representations of processes as networks of reactions and network motifs [18, 33–36]; although these coarse representations often do not account for kinetics or enzyme proximity, they have helped to establish bounds on the function of strongly-coupled reaction networks [37]. More sophisticated models accounting for enzyme size [38–41], charge[42, 43] and co-distribution [44] are based on ordinary and partial differential equation formalisms that implicitly capture these effects. Recent ordinary differential equations (ODE) approaches that implicitly consider the distributions of finite-sized enzymes include a mean field theory from Rao *et al*and[45]. These models provided strong quantitative insights into the efficiency of catalytic processes [46] and limits on efficiency gains for sequentially-coupled enzymes [45], but only implicitly account for geometrical and physiochemical factors. Explicit consideration of those factors for coupled enzyme processes generally utilize partial differential equations or particle-based solutions, which have afforded descriptions of how neighboring reactive enzymes [29, 47, 48], feedback inhibition, [29, 46], protein geometry and electrostatic interactions [19, 21, 49, 50] contribute to enzyme activity. Still, a systematic study of sequential reaction phenomena under physiological variations in solvent ionic strength and intracellular confinement is needed for probing how nucleotide pools are regulated by sequential NDAs in geometrically-constrained intercellular junctions.

Our key objective was therefore to quantify how cellular and organelle membranes tune NDA sequential enzyme activity and local nucleotide pools. This study was investigated in a model, biomimetic material for which the material porosity and surface composition could be controlled. Our approach uses a finite-element based partial-differential equation model developed in [51], for which we introduced explicit enzymes [23] to quantify how conditions such as lipid charge, ionic strength, porosity tune the efficiency of protein functions that utilize diffusing molecular substrates. Our key findings are that NDA colocalization and their charge complementarity with substrates can offset reduced reaction rates owing to their confinement in nanoscale volumes; moreover, tuning of pore surface properties further improve nucleotidase reaction efficiency. Through precise control of co-localization, the ratios of signal can be controlled. In this regard, NDA activity is strongly influenced by its environment, which can lead to reaction kinetics that differ relative to *in vitro* measurements.

## 3 Results

### 3.1 Overview

We have developed a computational model of sequential nucleotidase reactions confined to a narrow space between apposed cells. The reactions are activated by ATP diffusing from neighboring cells. This geometry consists of a pore with nanometer-scale radius spanning between two reservoirs to emulate a femtoliter-scale volume confined between two coupled cells. We impose an ATP gradient oriented parallel to a micrometer-length pore to reflect nucleotide diffusion into the pore. We summarize the conditions run in Table S1.

Within the pore we consider two sequential, CD39- and CD73-catalyzed ATP and AMP hydrolysis reactions that are in steady-state. With this model, we examine how enzyme co-localization, ‘tethering’ the enzymes to the pore wall, and charges on the enzyme and pore surfaces shape enzyme kinetics within the idealized pore volume. Although we assume that the enzymes are spherical with uniform reactivity and charge, we have found that such representations are reasonable approximations of structurally-detailed, non-uniformly charged proteins we have examined in other studies [52, 53], but are considerably less computationally expensive to evaluate.

### 3.2 Effects of molecular pore confinement on enzymatic activity

Coupled enzyme reactions have been widely studied in a variety of contexts, including isolated globular enzymes and along surfaces. Here we extend these approaches to examine nucleotide hydrolysis reaction kinetics for NDA enzymes embedded within nanoscale gaps between cells, which we emulated with nanopores of varying radii (see Fig. 1). These pores are representative of small, well-contained extracellular volumes, such as junctions formed between adjacent cells or synapsing neurons. We vary the relative distance between enzymes to investigate how co-localization impacts reaction efficiency, as well as their distance to the pore as to simulate surface-tethered versus freely-floating enzymes. In such geometries, substrate access to the enzyme is restricted to a narrow volume, which is expected to decrease the diffusion-limited reaction rate. To quantify the dependence of enzyme reactivity on nanopore volume, we numerically solved Equation 12 for substrate species ATP, AMP and Ado subject to the boundary conditions defined in Table 1. We defined the reaction efficiency as the ratio of the substrate Ado production rate coefficient, *k_prod,C_* over the substrate ATP association rate coefficient, *k_on,A_*. Since the association rate of AMP is equal to the Ado production rate, the reaction efficiency highlights how the reactivity of AMP is shaped by the system configuration independent of the ATP reaction rate. This ratio is identical to the ratio of the association rate coefficient of AMP to the production rate coefficient of AMP. We further note that this ratio of rate coefficients is identical to the ratio of particle flux values. We will use this identity to simplify the calculations, as the flux values are a more direct result of the numerical simulations.

**Figure 1:**
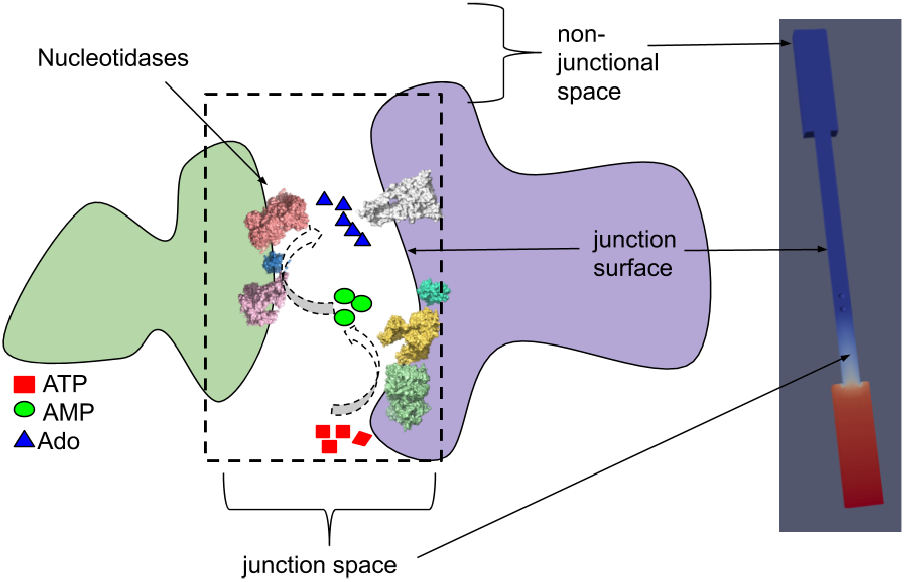
Left) Schematic of the synapse-like junctional space formed between the membranes of adjacent cells. Nucleotidases confined within the junctional space hydrolyze ATP into AMP and Ado. Right) A model geometry based on the schematic, for which the reservoirs correspond to the non-junctional space. The spatial and electrostatic configuration of the mock synapse influence the reactivity of confined nucleotidases CD39 and CD78..

**Table 1:** Concentration boundary conditions for the nanoporous system. Boundary conditions on Γ_*P*_ vary. Here, 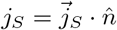.

| Boundary Surface | $\Gamma_{CD39}$ | $\Gamma_{CD73}$ | $\Gamma_R$ | $\Gamma_L$ |
| --- | --- | --- | --- | --- |
| Substrate ATP | $c_{ATP} = 0$ | $j_{ATP} = 0$ | $c_{ATP} = C_0$ | $c_{ATP} = 0$ |
| Substrate ADP | $j_{AMP} = -j_{ATP}$ | $c_{AMP} = 0$ | $c_{AMP} = 0$ | $c_{AMP} = 0$ |
| Substrate Ado | $j_{Ado} = 0$ | $j_{Ado} = -j_{AMP}$ | $c_{Ado} = 0$ | $c_{Ado} = 0$ |

We first validate our model against an analytical solution for the diffusion limited reaction rate coefficient on a uniformly reactive sphere. [54] Here, the association rate, *k_on_*, for the reactive sphere embedded in an infinite domain is given by

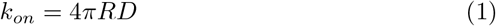

where *R* is the radius of the enzyme and *D* is the substrate diffusion coefficient. For the purpose of validation, we evaluate this rate at the sphere (0.5 nm radius) by assuming a uniform concentration for ATP (1.0 mM) at both the reservoir and pore (11.5 nm diameter). We will later assume no-flux (reflective) boundary conditions for the concentration along the pore to emulate a typical non-reactive pore boundary. Under the aforementioned conditions, we numerically estimated a rate of k_*on,ATP*_=3.611 × 10^−3^ nm^3^ ns^−1^, which is within 5% of the analytical estimate of *k_smol,bulk_*=3.768 × 10^−3^ nm^3^ ns^−1^. The minor discrepancy can be attributed to the nonspherical domain used for the numerical simulation, whereas a radially-symmetric domain is assumed in Eq. 1. We hereafter refer to this as the ‘bulk’ configuration, for which the enzyme concentration corresponds to roughly 0.19 mM. To establish a frame of reference for our results using reflective pore boundaries (see Table 1), in Fig. 2 we demonstrate a numerical prediction of *k_on,ATP_*=1.151 × 10^−3^ nm^3^ ns^−1^, which is within 1.8% of the analytical result, *k_smol,local_*=1.13 × 10^−3^ nm^3^ ns^−1^, for which we used the concentration. For this estimate, we average the bulk ATP concentration of the sphere centered within the pore. Both approaches confirm the reliability of the model for reproducing analytic results for diffusion-limited association reactions.

**Figure 2:**
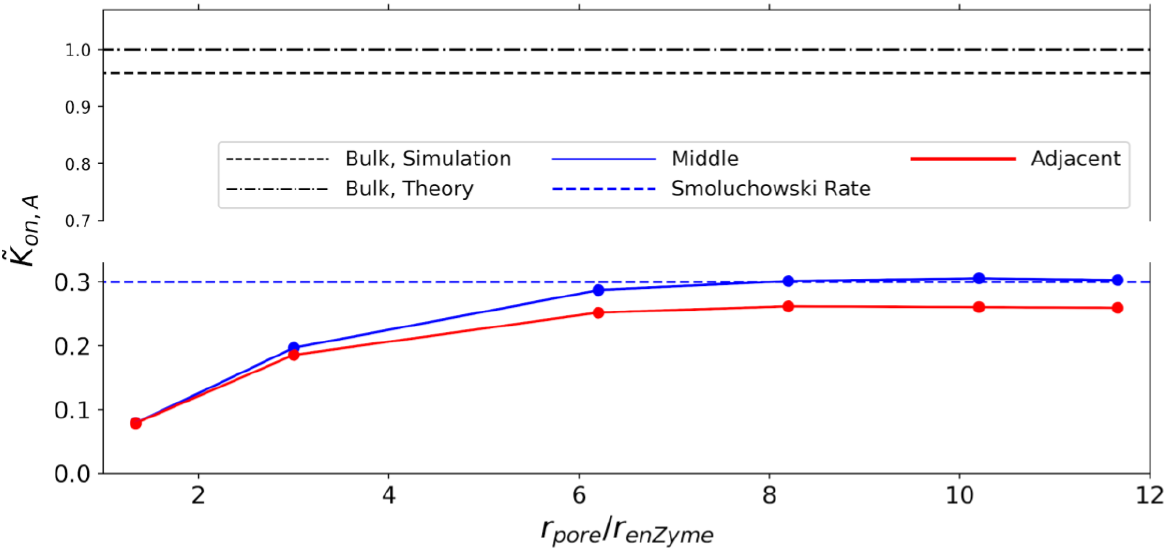
Effects of confinement and proximity. Normalized reaction rate coefficients of ATP to CD39, including comparison of simulation results to the analytical value for bulk conditions based on the Kimball-Collins relation. The vertical axis is broken to emphasize the dependence on pore radius..

Using our validated model we investigated how the reaction of substrate ATP on CD39 is influenced by confinement within a nanoscale channel. These simulations were conducted assuming a constant concentration gradient along the dominant axis of the channel. In Fig. 2) we present normalized association rates for ATP with enzyme CD39, 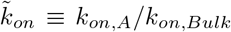, subject to a constant enzyme radius (*r_E_* = 1.0 nm) and varied pore diameters (*r_E_ ≈ r_p_* with *r_p_* =1.3 nm, *r_E_ < r_p_* with *r_p_* =3.0nm, and *r_E_ ≪ r_p_* with *r_p_* =5.5nm). Confinement of the enzyme to the pore reduced the reaction rate coefficient by roughly 70% relative to the corresponding rate in bulk 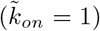. This can be qualitatively rationalized by the concentration profiles manifest in the channel (see Fig. S1). The concentration profile decreases from A=6.0 × 10^−4^ nm^−3^ at the right-hand side reservoir (Γ_*R*_) and approaches zero at the left-hand side reservoir (Γ_*L*_). As the pore radius decreases from *r_p_ ≫ r_E_* to *r_p_ > r_E_*, the concentration of ATP within the pore decreased relative to the reservoir. Hence, pore confinement in essence reduces the substrate concentration at the enzyme surface, which culminates in a reduced k*_on,ATP_*.

We additionally varied the proximity of the enzyme to the pore surface. This variation serves as a proxy for probing the reactivity of enzymes that are essentially free floating within the pore interior versus immobilized to the pore surface. The reactivity of CD39 was additionally reduced, albeit negligibly, as CD39 was localized to the pore surface. This can be rationalized by noting the similarity between the time-independent diffusion equation and the Laplace equation commonly used in electrostatics (see Eq. 15 with *κ* = 0.).

The total electric flux is dependent on the capacitance; as the sphere approaches an insulator wall (*J · n* = 0), the capacitance decreases [55], which decreases the total electric flux. The total electric flux is the electrostatic equivalent of the concentration flux in substrate diffusion, hence, the numerically-estimated k_*on,ATP*_ values were smaller for immobilized enzymes relative to those far from the surface. Altogether, these results demonstrate that restricting the diffusion of ATP within the pore and to a slightly greater extent, near the pore wall, suppress k_*on,ATP*_ relative the the bulk.

Reduction of k_*on,ATP*_ through enzyme confinement of enzymes is expected to subsequently suppress production rates for AMP and Ado. However, co-localization of enzymes within ‘nano reactors’ is a common approach to tune production rates of desired chemical products [44, 46, 56]. We therefore introduced a second enzyme, CD73, into the pore and simulated the steady state reactions *ATP → AMP* at cD39 and *AMP → Ado* at CD73. In Fig. 3) we first report Ado production rate coefficients, k_*prod,Ado*_, as a function of enzyme separation and for non-reactive (reflective) and reactive (absorbing) pore surfaces. These values are normalized with respect to the k_*prod,Ado*_ value obtained for *r_p_ ≫ r_E_* and maximal enzyme separation (*d_CD39,CD73_*=1). The normalized kprod,Ado rate coefficients are negligibly impacted as enzyme separation is reduced within a boundary that is nonreactive with AMP (e.g. reflective); this indicates that enzyme colocalization has negligible impact on k_*prod,Ado*_ if AMP does not interact with the pore boundary (e.g. reflective). If absorbing conditions are assumed, that is AMP is depleted at the pore, then there is a kinetic advantage to enzyme colocalization; this is demonstrated by the increased rates with reduced 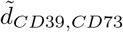 in Fig. 3) that are accentuated for increasing pore sizes. Hence, as the domain approaches the bulk-like system where AMP can escape from the reaction complex, the advantage of enzyme proximity becomes apparent and is consistent with recent studies of enzyme co-localization [19, 45, 46]. In summary, the nature of the intermediate, AMP, interactions with the surface appear to determine the relative advantage of enzyme colocalization in closed, nanoscale domains. For example, in a scenario where enzymes auxiliary to NDAs are depleting nucleotides, NDA enzymes would benefit from co-localization.

**Figure 3:**
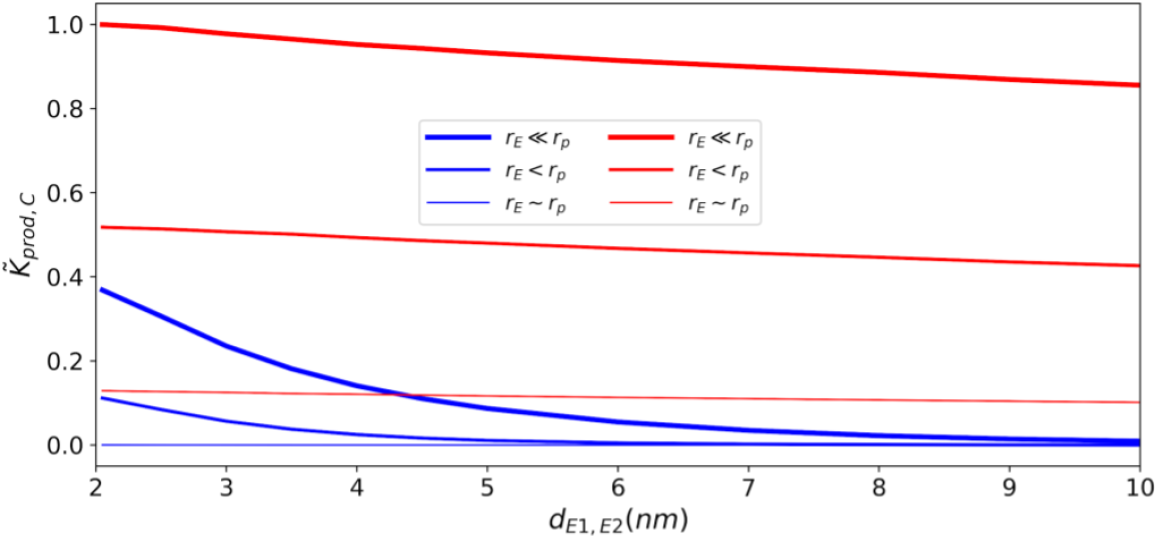
Effects of confinement and proximity. Normalized reaction rate coefficient for the second enzyme at different pore sizes and enzyme proximities. Blue lines are for absorbing boundary conditions to emulate an open, bulklike configuration, and red lines are for reflecting boundary conditions. 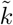 is normalized with respect to the maximal Kprod value at min d max rp..

To delineate the effects of pore confinement and enzyme colocalization on k_*on,AMP*_ and k_*prod,Ado*_ independent of k_*on,ATP*_, we report in Fig. 4) the Ado production efficiency, k_*eff*_, which we define as k_*eff*_≡k_*prod,Ado*_/k_*on,ATP*_. The efficiencies reported for co-localized enzymes (left panel) are consistently higher than those for separated enzymes (right panel), with the largest increases demonstrated for reactive pore boundaries (blue) or open (red, bulk) configurations. Hence, under circumstances that permit intermediates to diffuse away from or compete with the reactive centers, there is a clear advantage to colocalization, akin to findings in [45]. However, in the absence of substrate interactions with the pore surface, confinement leads to higher overall efficiencies that monotonically decrease with increasing pore radius, but have little dependence on enzyme proximity.

**Figure 4:**
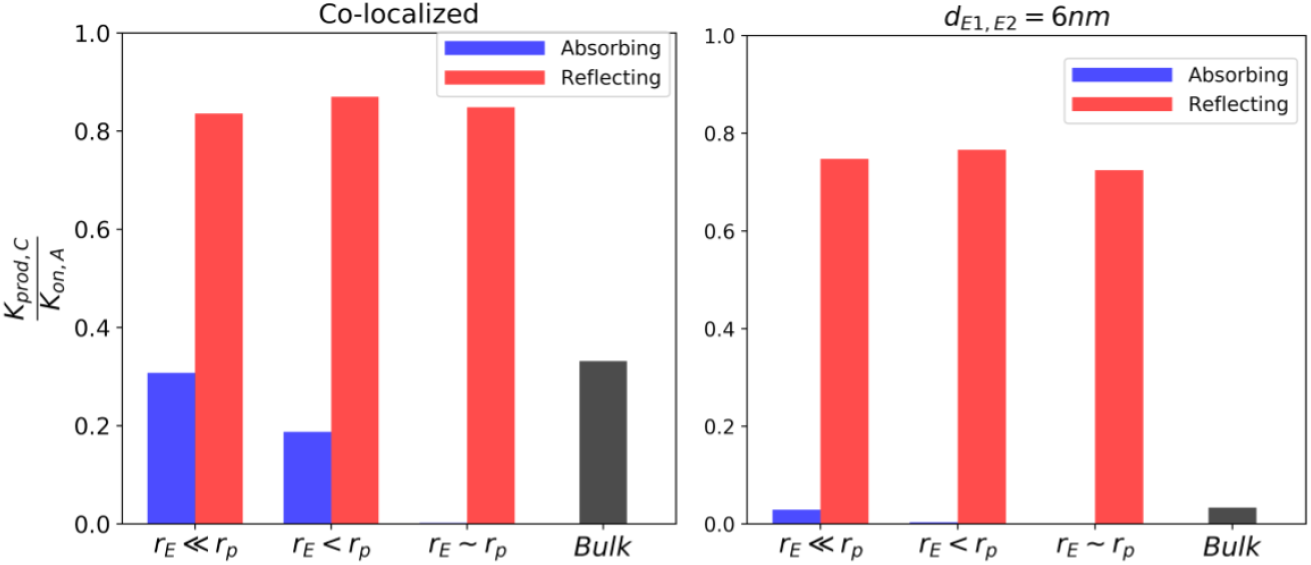
Effects of confinement and proximity. Efficiency of reactivity of second enzyme respect to first enzyme. Left panel: enzymes co-localized. Right panel: enzymes separated by 6 nm. Efficiencies for the absorbing case approach 0 as *r_p_ → r_E_*.

**Figure 5:**
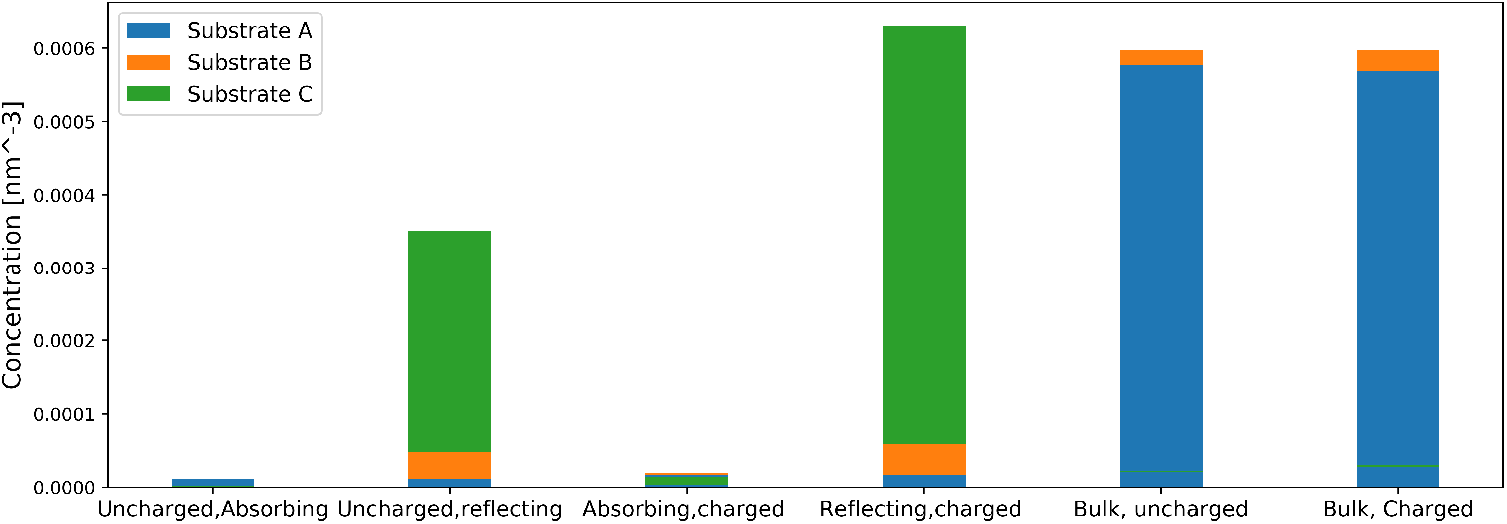
ATP, AMP and Ado concentrations at midpoint between enzymes for different pore wall boundary conditions. CD39 and CD73 are separated by a distance of 6.0 nm..

### 3.3 Effects of surface charge on reaction rate coefficient

In the previous section we demonstrated that confinement of enzymes like the ectonucleotidases to nano-scale extracellular domains suppresses the overall reaction rate coefficient of uncharged substrates. Co-localization of the reactive centers mitigated this reduction to a modest extent. Naturally, the adenosine substrates are charged, with ATP having the most negative charge and AMP the least. Hence, their concentrations and diffusion rates are expected to be sensitive to the charge configuration of their binding partners and the surrounding lipid bilayer environment. It is well-known, for instance, that many enzymes have evolved to exploit electrostatic interactions to accelerate substrate binding [42]. Further, there is strong evidence that local ionic concentration near charged membranes yield concentrations of Ca^2+^ and Na^+^ that deviate significantly from the bulk [57, 58]. We therefore expanded the approach in the previous section to consider competing or complementary effects of electrostatic interactions in coupled enzyme kinetics.

In this section, we model the contribution of electrostatic interactions between substrates and their environment using the Smoluchowski electro-diffusion equation (see Eq. 8), for which the electrostatic potential was modeled using the linearized Poisson-Boltzmann equation (see Eq. 15). We first validate our implementation under dilute solvent conditions (*κ* → 0) and assume that the pore and CD73 are uncharged. The association rate for a substrate with charge *q_A_* with a spherically-symmetric enzyme of radius, *r_E_* and charge *q_E_* can be analytically determined [19]:

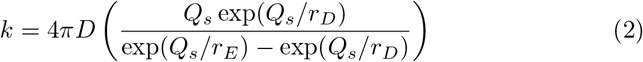

with *Q_S_* ≡ *q_A_q_E_*λ_*B*_, where *r_D_* is the radius of the domain within which the reaction is confined and λ_*B*_ is the Bjerrum length. Accordingly, we demonstrate in Fig. S2 that as *κ* → 0 (log(*κ*) → −∞) the numerically-predicted k_*on,ATP*_ rates for *z_ATP_* =−1 and Φ_*CD*39_ = 25 mV approach the analytical estimate within 6% percent, and thus reasonably validate the electrostatic model.

Using the validated electro-diffusion model, we first examined changes in CD39 reactivity by confinement within an electrically-neutral pore, subject to electrostatic interactions between a negatively charged substrate A, positively charged CD39 and variably charged CD73. We assumed surface potentials of ±19.2 mV for the enzymes based on *ζ* potentials measured for proteinaceous solutions by Salgn *et alet al* [59]. Further, although adenosine metabolite charges vary from −4 to 0, and because Ado is commonly chelated by Mg^2+^ [60], we used charges of −2, −1 and 0 for ATP, AMP and Ado, respectively, to exemplify effects on reactivity. Under these conditions, k_*on,ATP*_ decreases as the pore radius to enzyme radius decreases as was observed for the neutral system (see Fig. 6). Importantly, k_*on,ATP*_ for the charged system does assume a higher rate coefficient than the neutral system. Hence, the electrostatic interactions in essence counterbalance the reduction in reaction rate coefficients due to confinement. Moreover, NDAs complementary charge with nucleotides exploits fast association. Additionally, consistent with [19], when CD73 and CD39 assume the same charge complementary to the intermediate, increased k_*on,ATP*_ rate coefficients result, owing to the attraction between enzymes and the substrate.

**Figure 6:**
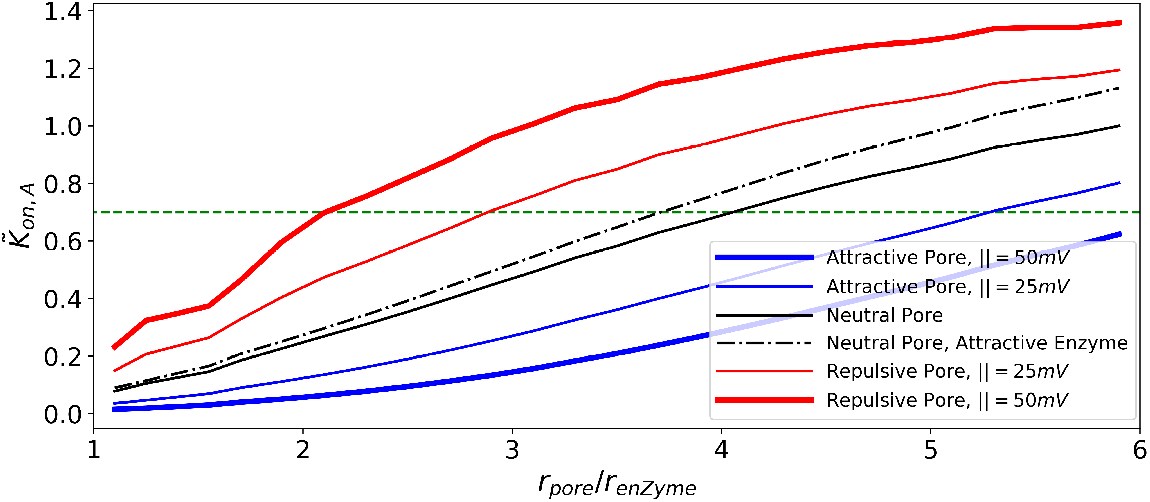
Effects of charge. Effect of charge sign composition on reactivity, given *z_ATP_* =−1, *z_AMP_*=+1, *z_Ado_*=0 and Φ_*CD*39_ > 0 Effect of pore electrical potential on reactivity of enzyme for different sizes of the pore. We define an effective pore radius for charged pores, to be obtained from the green dashed line. For relatively small pores, the effective pore radius is larger for attractive pores, and smaller for repulsive pores. For relatively larger pores, competition between the pore wall and the enzyme becomes more significant, leading to a decline in the reactivity of attractive pores (see Fig. S3 in which with decreasing the size of enzyme we allow larger relative sizes of pore to enzyme radius)..

We next imposed a negative electric potential on the pore surface and present the resulting reaction rate coefficients (red in Fig. 7). We chose surface charge densities consistent with biological membranes reported by surface conductivity microscopy such as DPTAP=15.1 mCm^−2^, DPPE=5.3mCm^−2^ and DPPG=−44.0 mC m^−2^ for positively charged, zwitterionic and negative charged lipid bilayers, respectively [61]. We first examine effects of the pore electric potential, Φ_*pore*_, on reaction kinetics, assuming CD73 is uncharged. In Fig. 7 we demonstrate that in general k_*on,ATP*_ monotonically decreases regardless of the membrane charge. In the event that the pore interactions with substrate ATP are repulsive (Φ_*p*_ < 0), the reaction rate coefficient decreases at a faster rate. However, in certain regimes the charge complementarity of the pore surface was found to greatly accelerate k_*on,ATP*_ relative to the neutral pore, whereas a repulsive pore (Φ_*pore*_ < 0, blue) attenuated k_*on,ATP*_ by roughly 17% or 3.77 × 10^−1^ nm^3^ ns^−1^ (see Fig. 6) for *r_p_* ∼ 6*r_E_*). We attribute the enhanced reaction rate coefficient for the positively charged membrane to the elevated concentration adjacent to the membrane relative to the uncharged membrane, which effectively raised its average concentration within the pore (). That is, the complementary charged pore surface drew ATP into the pore interior and thereby facilitated the reaction on CD39. Hence, the charge of the pore surface stemming from different phospholipid compositions can strongly influence k_*on,ATP*_, and in turn control AMP productions ATP degradation via CD39.

**Figure 7:**
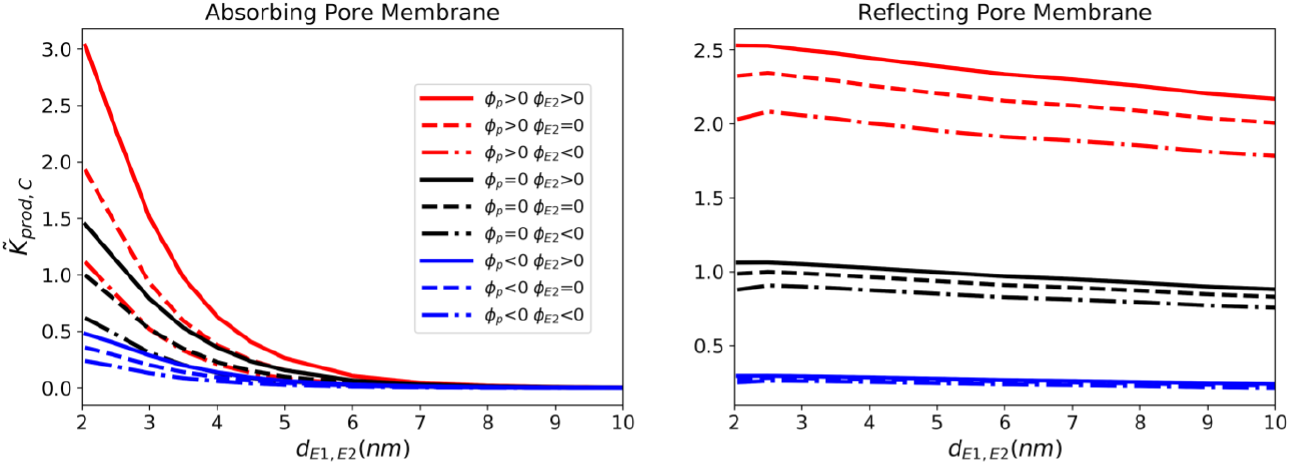
Effects of charge. Effect of charge sign composition on reactivity, given *z_ATP_* =−2, *z_AMP_* =−1, *z_Ado_*=0 and Φ_*CD*39_ > 0 Normalized reaction rate coefficient for production of Ado as a function of distance between enzymes..

Interestingly, attractive pore/ATP interactions initially accelerate k_*on,ATP*_ as the pore diameter is reduced, whereafter the rate declines. We find the maximal acceleration is achieved when the pore size is roughly six-fold higher than the enzyme radius (see Fig. S3). This maximum is dependent on the wall potential amplitude, namely as the attractive wall potential amplitude increases, the maxima shift to smaller pore/enzyme size ratios. In previous studies [51, 52, 62], it has been demonstrated that weakly attractive interactions with pore boundaries can enhance diffusion and ion conductivities, which is consistent with the initial increase in k_*on,ATP*_ in our model. However, this acceleration in diffusion due to attractive interactions is eventually outweighed by the hindrance of diffusion as the pore is narrowed. Additionally, although diffusion is likely accelerated, the amount of substrate able to interact with the target is reduced, as we saw for uncharged cases. Hence, attractive pore potentials served to co-localize substrates near the pore wall and therefore offset the reduced reaction volume that would otherwise decrease the overall reaction rate coefficient (see Fig. S7), as might be expected for ATP with positively charged phospholipids.

We initially anticipated that positioning the protein directly adjacent to the membrane would improve the reactivity relative to the pore center. Further, we observe that k_*on,ATP*_ can be amplified when the enzymes are tethered to the pore surface under specific conditions, namely wide pores and strong attraction, but this advantage is generally minor and thus of limited consequence to NDAs (Fig. S6). We found little difference for k_*on,ATP*_ at modest (< |25| mV) pore potentials for far in Fig. S7). This appears to be consequence of reduced access to the enzyme as it approaches the wall, which counterbalances the increased the concentration of ATP near the surface due to attractive electrostatic infractions.

Electrostatic enhancement of k_*on,ATP*_ is generally expected to promote k_*on,AMP*_ and k_*prod,Ado*_, thus we examined the extents to which the intermediate species’ charge and enzyme proximity control k_*eff*_. We present results assuming a positively-charged intermediate (*z_AMP_*=1) to emphasize the electrostatic control of the sequential reactions. Consistent with findings from the neutral system and our previous studies of sequential enzyme channeling [19], we find that k_*prod,Ado*_ increases as the enzymes are brought into close proximity 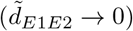. As observed in the preceding section, the absorbing pore boundaries show the greatest sensitivity to enzyme distance, with favorable AMP /CD73 electrostatic interactions for Φ_*CD*73_ < 0 yielding faster k_*prod,Ado*_ reaction rate coefficients relative to neutral CD73, and conversely slower rates for positively-charged CD73. The enhancement in the former case reflects both the electrostatic attraction of substrate AMP toward CD73, while ATP is electrostatically repelled toward CD39, which culminate in increased k_*on,AMP*_ and k_*prod,Ado*_, respectively. In the latter case, the positively charged CD73 repels AMP and attracts the negatively-charged A, which in effect competes with the reaction of ATP on CD39. Hence, the enzymes’ charge complementarity with their respective substrates enhances the overall reaction rate coefficient relative to uncharged systems and thereby offsets the suppressed rate due to confinement within the pore. In this regard, co-localization of the nucleotidases can ensure reasonable reaction efficiency despite confinement in pore. Further, these results confirm trends identified in [19, 45, 46] that enzyme colocalization can support higher overall reaction rate coefficients under specific conditions, albeit here we consider such effects in the context of confinement to the pore.

Attractive interactions between substrate ATP and a positively-charged surface lead to the greatest overall enhancements of k_*prod,Ado*_. There is, however, a limit to this acceleration, if the attractive interactions between the pore and substrate are stronger than the target enzyme, as we demonstrate with a reduction in k_*on,ATP*_ demonstrated in Fig. S6. It is interesting that attractive A/pore interactions dominate the sequential enzyme kinetics, given the likely repulsion of cationic B species from the reactive centers, which could suppress the overall reaction efficiency. To assess the extent to which AMP /pore interactions contribution to k_*prod,Ado*_, we examined the k_*eff*_ ratios for the nonreactive pore boundaries. Based on the efficiencies reported in Fig. 8, most of the increased production rate can be attributed to k_*on,ATP*_, as the efficiencies were largely constant across the various charge configurations. However, efficiencies were generally greater for co-localized enzymes versus separated configurations, regardless of the pore charge For negatively-charged pores (Φ_*pore*_ < 0), the reaction efficiency was enhanced for all configurations, except for colocalized enzymes directly adjacent to the pore. In analogy to the effects of the pore charge on ATP, the increased efficiency for positively-charged pores can be rationalized by the pooling of anionic AMP within the pore, which increases k_*on,AMP*_. This concurs with our findings of enhanced reaction efficiency in dihydrofolate reductase-thymidylate synthase (DHFR-TS) [21] owing to its complementary surface charge to the dihydrofolate intermediate intermediate, in contrast to a neutral or electrically-repulsive surface. For the co-localized cases at the membrane surface, it is likely that the membrane significantly competed with the binding of B at CD73, which led to a reduced reaction rate coefficient. Overall, these results suggest that favorable A/membrane interactions largely control the absolute k_*prod,Ado*_ rate, with co-localization typically further enhancing k_*prod,Ado*_ and k_*eff*_. The contributions of B/pore interactions play a comparatively smaller role, perhaps given the limited extent to which B permeated the pore relative to A.

**Figure 8:**
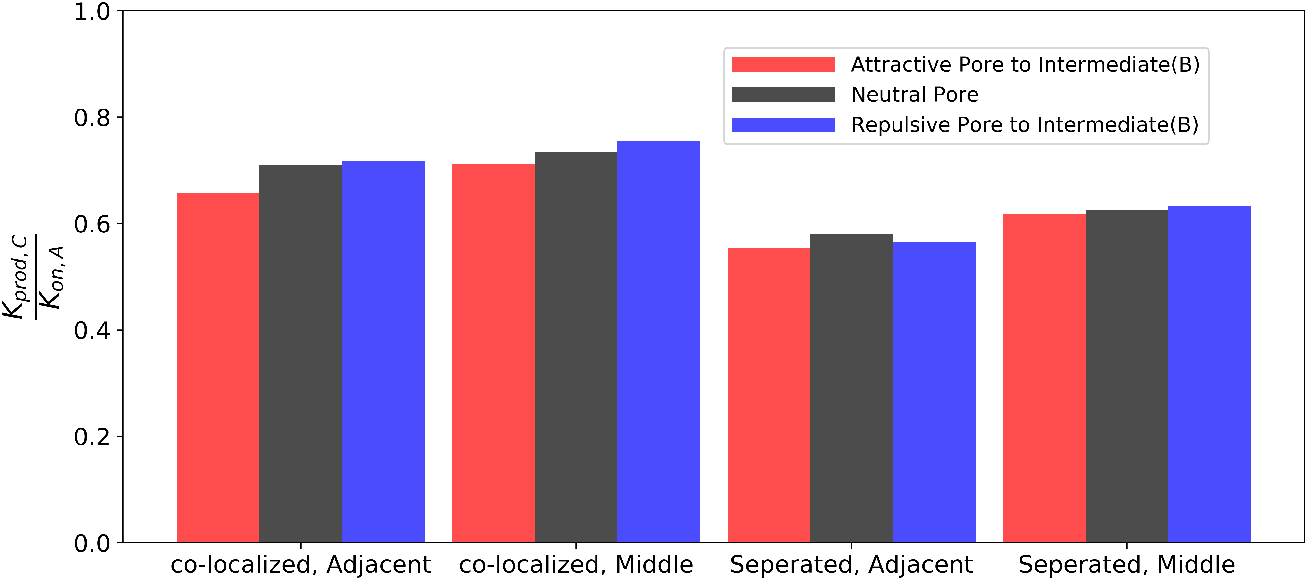
Effects of charge. Efficiency of sequential enzymes for different pore wall electric potentials, given *z_ATP_* =−2, *z_AMP_* =−1, *z_Ado_*=0 and Φ_*CD*39_ > 0. Corresponding values for k_*on,ATP*_ are provided in Fig. S7.

In the previous section, we highlighted reaction rate coefficients and efficiencies without accounting for electrostatic screening by common electrolytes. To model physiological conditions characterized by roughly 100 millimolar monovalent ion concentrations, we solved the linearized Poisson-Boltzmann equation, assuming Debye lengths on the order of 1 nm. This Debye length signifies that electrostatic interactions are significantly screened, which will in turn modulate reaction kinetics. To assess effects on k_*on,ATP*_, it is helpful to compare rates as a function of (1 + *aκ*), where a is the enzyme radius and *κ* is the inverse Debye length (see Fig. 9). This functional form is motivated by the relationship

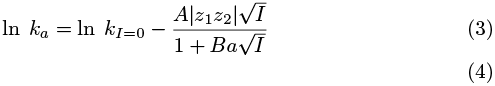

where I is the ionic strength, while A and B are generally fitting parameters introduced by Schreiber *et al* [63]. This latter term stems from the Debye-Huckel treatment of electrolyte solutions.

**Figure 9:**
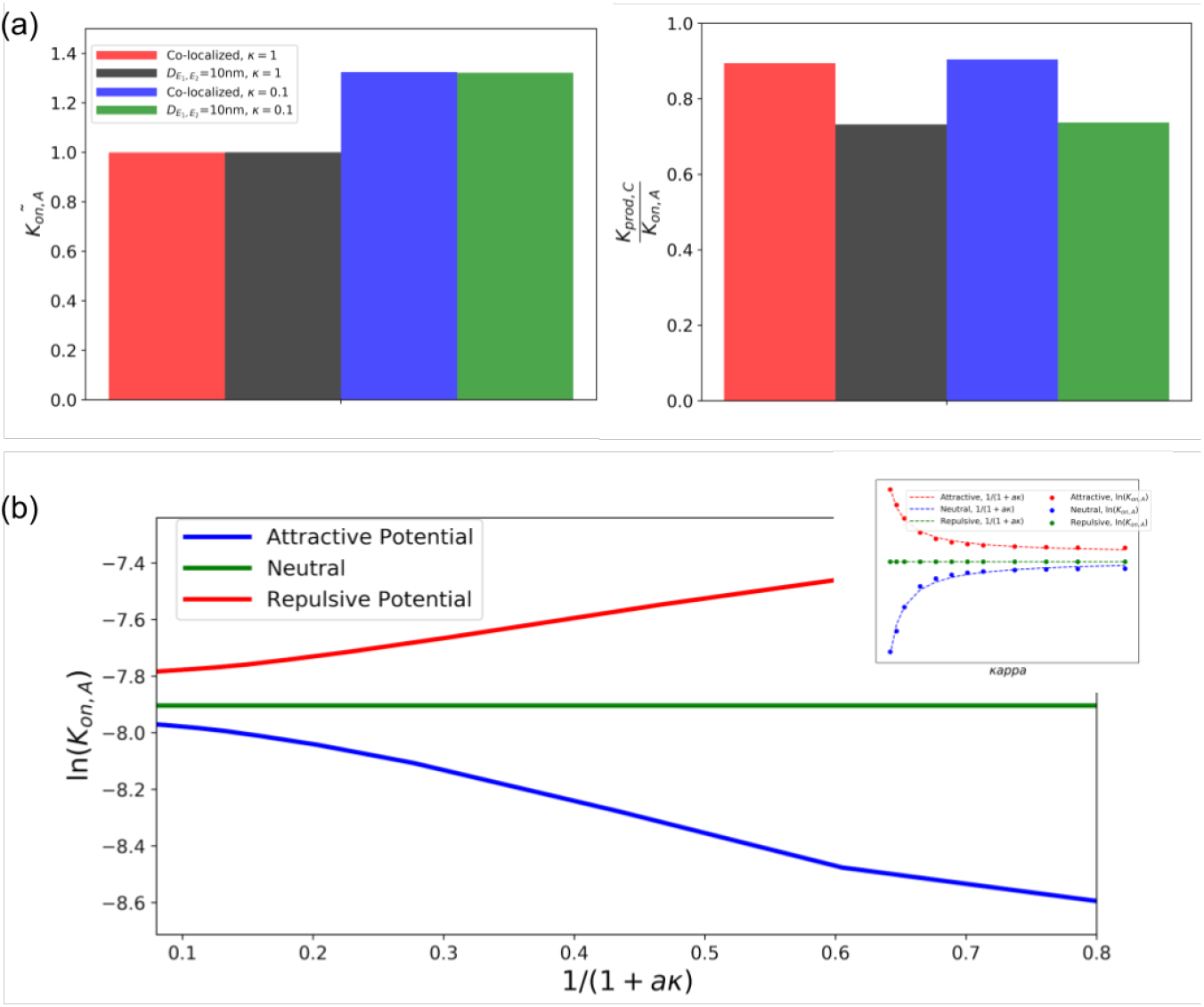
Effect of Debye Length on reactivity of sequential enzymes: a) Efficiency and first enzyme reactivity for large and small Debye lengths for co-localized and separated enzymes for Φ_*CD*39_ > 0, Φ_*p*_ > 0 and Φ_*CD*73_ < 0 *z_ATP_*=−1 and *z_AMP_*=1 b)Relationship between ln(*κon*) and (1 + *aκ*)^−1^ for three different pore electrostatic potentials. Subpanel shows fitting of curve to simulation data points.

In Fig. 9a), we demonstrate the scaling of ln[*k_on,A_*] with respect to (1 + *κa*)^−1^ → 1, assuming either attractive or repulsive substrate interactions with CD39 and pore. For attractive interactions, the maximum rate enhancement relative to an electrically-neutral reference system is found under dilute conditions signified by *κ* → 0 or equivalently, (1 + *κa*)^−1^ → 1. At high ionic strength (*κ* → ∞), the rates approach those of the neutral system. By accounting for electrolyte screening, reaction rate coefficients are somewhat attenuated, which will in turn depress k_*prod,Ado*_ and potentially k_*eff*_. Accordingly, we reevaluated the effects of enzyme co-localization and pore proximity on k_*prod,Ado*_ and k_*eff*_, subject to the 150 mM KCl background electrolyte. As anticipated, k_*prod,Ado*_ generally scales proportionally to k_*on,ATP*_ as a result of A/pore interactions largely setting the overall reaction rate coefficient relative to the intermediate. This is further evident in the negligible impact of ionic strength on k_*eff*_. We additionally found that the trends in k_*eff*_ and k_*prod,Ado*_ relative to enzyme co-localization and pore proximity did not significantly deviate from those reported in electrolyte-free conditions and are therefore not reported here. Overall, while physiological ionic strength conditions modestly impact reaction kinetics relative to dilute conditions, the changes are fairly insignificant and also insensitive to moderate changes in ionic strength typical in physiological systems. In other words, fluctuations in signaling ion concentrations are not likely to significantly modulate NDA behavior.

## 4 Discussion

### 4.1 Summary

In this study, we probed how nucleotide pools are controlled by ecto-nucleotidase (NDA) and nucleotidase enzymes within narrow junctions formed between adjacent cells, similar to synaptic junctions comprising neurons. Here we used numerical solutions of steady-state reaction-diffusion models of coupled nucleotidases that were recently studied in cystic fibrosis but assuming spatially uniform behavior [31]. These simulations were performed under physiological conditions that included confinement to narrow junctions bordered by cell plasma membrane and long-range, ionic strength-mediated electrostatic interactions. This three-dimensional approach resolved important factors governing the fast reaction rates of NDAs in biological systems, that were not readily apparent in prior numerical approaches based on ordinary (spatially-independent) differential equations. Our key findings were

- that independent of NDA activity, nucleotide distributions within confined extracellular junctions can significantly differ relative to open, bulk-like configurations
- that NDAsteady-state reaction rates are generally smaller when localized to small junctions, but reaction efficiency can improve by co-localizing coupled enzymes, and
- that these reaction rates can be substantially accelerated when NDA and plasma membrane adopt charges complementary to reacting substrates, especially when the membrane attracts the relevant substrate.

Adenosine nucleotides encompass a set of small, polar molecules that are critical for cellular signaling and metabolism [14]. These nucleotides are generated or regulated by diverse processes, including secretion from neighboring cells in tissue [64], as products of membrane-bound F1-ATPases [65], transport via ectopic adenine nucleotide translocases [66]or hydrolyzed by ecto-nucleotidase (NDA). For many cellular systems, these processes occur within femtoliter-scale [11, 67] regions between neighboring cells, such as those characteristic of neuronal synapses [68]. Here, co-localization of NDAs including CD39 with purinergic receptors within caveolae [69] or with extracellular ATP release sites on astrocytes [70] can give rise to ‘compartmented’ nucleotide pools [71] that can strongly influence nucleotide-dependent signaling. The thermodynamics and kinetics of molecular signaling in such compartments can differ considerably from analogous processes in bulk solutions or in vitro. Myriad factors contribute to these differences, including smaller compartment volumes that strongly amplify substrate concentration gradients [30, 72], the presence of ‘crowders’ comprising other small molecules, protein or nucleic acid that generally impede diffusion [28, 47, 73, 74], as well as rate enhancements typically exhibited for closely-apposed enzymes [29, 42, 45, 50, 75] or those adopting electrostatic fields complementary to a reacting species [38]. The relative contribution of these factors to coupled NDA activity in a given multi-cellular domain has not been examined previously and could ultimately determine the relative distribution of nucleotides. This balance of nucleotide concentrations determine the extent to which membrane bound, nucleotide receptors, ATPases and translocases are activated, that in turn can shape diverse cellular processes, including migration [76] and cytokine release [77].

To systematically probe the potential for these factors to impact NDA activity and nucleotide distributions, we examined a sequential adenosine nucleotide hydrolysis process as a model system for characterizing ecto-nucleotidase (NDA) activity within synapse-like domains. We performed steady-state reaction-diffusion simulations of two NDA enzymes that sequentially catalyze the hydrolysis processes *ATP/ADP* → *AMP* → *Ado*. Our system is modeled after the CD39 and CD73 NDAs, though we utilize two arbitrary, spherical representations that are selective for tri- and mono-phosphates, respectively, to generalize our results to other coupled nucleotidases. Additionally, although the fully deprotonated ATP anion assumes a charge of −4 [78], we assume in accordance with physiological systems that it is coordinated with magnesium [79] and thereby assumes a net negative charge of −2. To orient our results, we note that increasing k_*on,ATP*_ rates (reaction of ATP at CD39) result in ADP accumulation, while large k_*on,AMP*_ and k_*prod,Ado*_ correspond to high rates of ADP consumption and AMP production at CD73. Since the latter rates are generally proportional to k_*on,ATP*_, we report the reaction efficiency, k_*eff*_ =k_*prod,Ado*_/k_*on,ATP*_, to highlight contributions specific to ADP consumption. In this regard, high k_*eff*_ rates generally reflect significant consumption of extant ADP pools to form AMP. These rates were measured for CD39 and CD73 in bulk solution and confined within a nanometer-diameter pore, for which we varied the enzyme distributions, substrate/enzyme interactions, as well as the nanopore surface charge to emulate typical phospholipid bilayers. Based on these variations, we discuss how the relative nucleotide composition within confined domains is determined by physical attributes of the nanoscale compartment, how co-distribution of coupled hydrolysis reactions helps facilitate high reaction efficiency despite confinement to nanoscale, extracellular junctions, how membrane surface charge localizes substrates so as to accelerate reaction rates, indirectly modulate enzyme kinetics.

### 4.2 Nucleotide transport and distribution (pools) within crowded extracellular junctions

We first discuss how nucleotide diffusion rates and distribution are influenced by physical attributes of the confined junctional geometry, including restricted diffusional volumes and electrostatic interactions between substrates, reactive enzymes and charged membrane surfaces. Independent of NDA activity, the restricted volume of the junction relative to the surrounding substrate reservoir, as well as the surface charge distribution within the junction, played key roles in shaping the nucleotide distribution. In our model, nucleotides entered the restrictive junctional domain from one of two reservoirs to emulate entrapment of species generated from an external source, such ATP released from nearby damaged cells or part of physiological processes including paracrinic release. In the absence of nucleotide/surface interactions, the diffusion rate of nucleotides through the junction decreases as its radius is reduced. This is easily rationalized by noting that the substrate flux through a cylinder normal to the nucleotide concentration gradient scaled proportionally to the cylinder’s cross-sectional area relative to the reservoir surface area [80]. The constriction of the substrate-accessible volume at the junction opening leads to a substantial reduction in the amount of substrate available to the enzyme within the pore compared to bulk conditions (see also [51]). For this reason, narrow junctions between cells are anticipated to limit nucleotide pools available to ATPases and ATP-gated receptors localized to extracellular junctions. Hence, estimates of ATP based on bulk (extracellular) measurements [79] are generally unrepresentative of the local ATP pools formed within the compact interstitia between cells. This deviation strongly justifies the use of localized measurements of nucleotides when probing receptor activity in neural synapses for instance via microelectrodes [10].

### 4.3 Contribution of NDAs to controlling nucleotide pools

A secondary focus of this computational study was to probe NDA dependent modulation of steady-state nucleotide concentrations relative to those determined by junction size and electrostatic charge alone. It is understood that NDAs rapidly degrade nucleotides released in synaptic junctions [81]; hence, pulsatile release of ATP from post-synaptic neurons is followed by transient, millisecond-scale upswings in the synaptic ATP concentration, owing to NDA degradation [10]. However, the femtoliter volume of such spaces [11] and intercell separations within several fold of the Debye length suggest that NDA activity and resulting nucleotide pools will be sensitive to NDA colocalization, strengths of substrate/enzyme electrostatic interactions and the junction volume. Firstly, in analogy to the reduced nucleotide concentration reported at the junction/reservoir boundary, we observed substantially lower k_*on,ATP*_ rates for junction-confined CD39 relative to the bulk configuration. This behavior is easily rationalized by the smaller cross-section of the pore relative to an open system, which both reduces the concentration of substrate at the enzyme surface, as well as the accessibility of reactive enzyme surface. Further, our predictions are consistent with classic theoretical studies relating the dynamic accessibility of gated protein active sites or substrate tunnels to observed enzyme activity [82], as demonstrated in acetylcholinesterase [83] and the PutA peripheral membrane flavoenzyme [84]. Since k_*prod,Ado*_ scales proportionally to k_*on,ATP*_, reduced NDA rates owing to confinement suggests that in vitro characterization of NDA activity in bulk media likely yield faster kinetics than would be expected for strongly confined systems; As a consequence, ATP pools within the junction were largely suppressed relative to those of the NDA-free system, while ADP and AMP were considerably larger. Based on these predictions, we anticipate that the degradation of adenosine phosphates to lower order molecules by ectonucleotidases proceeds more slowly in confined extracellular spaces relative to bulk conditions. Further, this reduction in reactivity is largely determined by the reaction rate of the first species, ATP.

In contrast to the consistent rate-limiting effect of enzyme confinement on k_*on,ATP*_, the efficiency of Ado production relative to bulk varied depending on the nature of substrate/membrane interactions. We investigated this dependency assuming reflective (non-interacting) and absorbing boundary conditions on the membrane; the latter is representative of nucleotide depletion by membrane-bound ATPases or translocases. We found that efficiency was maximized when the nucleotides did not significantly interact with the membrane (reflective). In this case, although CD39’s confinement to the pore limited its access to ATP, the membrane prevented intermediate diffusion away from CD73. This established a relatively high intermediate concentration within the junction that in turn increased k_*prod,Ado*_. In contrast, efficiency was strongly reduced when nucleotides were depleted at the surface (absorbing), as might be expected for significant nucleotide uptake by plasma membrane adenine nucleotide translocases [66]. As discussed in the next section, this reduced efficiency for absorbing membranes could be countered by co-localizing the two-enzymes to favor AMP ‘s reaction on CD73 relative to diffusing toward the membrane Ultimately, these findings suggest that nucleotide pools capable of activating targets such as ADP sensitive P2Y channels will be strongly regulated by the relative activity of proteins or transporters that reduce the di-phosphate concentration in the junction and thereby compete with CD39.

Numerous biochemical processes that involve diffusing reactants rely on close spatial coupling of enzymes to promote efficient signaling. Examples of enzyme co-localization include formation of macro-molecular complexes [85, 86], confinement in molecular ‘tunnels’ [87–89], the proximal reactive sites in the sulfate-activating complex [90], in addition to metabolic substrate channeling [91–93]. We had thus expected that co-localizing NDAs within junctions would improve reaction efficiency. However, we found that close spatial coupling was advantageous only when the junction membrane significantly interacted with the intermediate. Specifically, when the membrane either absorbed the intermediate or attracted the intermediate through attractive electrostatic interactions, smaller concentration gradients were evident at CD73 and thereby reduced k_*on,AMP*_. Co-localizing CD39 and CD73 minimized the intermediate’s access to the membrane and thus facilitated faster k_*on,AMP*_ rates than were evident at larger enzyme separations. This behavior is consistent with simulation studies by us and others [19, 45, 82] for open (bulk) systems whereby co-localization of sequential enzymes can enhance reaction rates. Based on our rationalization in the preceding paragraph, co-localization of CD39 and CD73 for a reflective membrane had minimal impact on the reaction efficiency, given that the intermediate had limited capacity to escape the reactive sites. Constructs including micelle- or viral capsid-based nanoreactors that house enzymes, or enzymes immobilized to linear or planar molecular assembles [56, 94] exhibit analogous increases in efficiency through mitigating loss of intermediates to open boundaries. These results therefore suggest that variations in NDA co-localization could provide a means to tune the relative composition of nucleotide pools within junctions, particularly for charged membranes or those with an abundance of proteins that compete for nucleotides.

A central contribution from our study is to confirm the significant role of electrostatics and intermediate channeling in facilitating coupled nucleotide hydrolysis reactions catalyzed by NDAs in nanoscale volumes. Secondarily, we demonstrate that tuning of the surface/enzyme and surface/substrate interactions can further optimize reaction rates. A third contribution of our study was to systematically characterize how electrostatic interactions influence enzyme kinetics under physiological conditions. It is clear from our simulation data that a significant membrane charge can redistribute the populations of charged substrates along the pore boundaries. Because diffusion-limited reaction rates scale proportional to the substrate concentration gradient at the enzyme active site, it was expected that membrane charge configurations that localized substrates to the pore and its surface would enhance the reaction rate.

From this standpoint, we can treat our predicted k_*on,ATP*_ values as readouts for the significance of local substrate concentrations to NDA activity, particularly in the context of electrostatic interactions. Such electrostatic interactions have been speculated to contribute to the formation of ‘micro-domains’ localized to the membrane surface, such as for Ca^2+^, Na^+^ and to a lesser extent [58, 72, 95, 96] following transient fluctuations in their concentrations. These microdomains are strongly implicated in modulating the ion-dependent activation of small proteins [97]. As an example, ATP has been suggested to assume concentrations several-fold higher than the bulk cytosol, based on modeling and ATPase enzyme assays [30, 93]. To the extent that the microdomains arise exclusively from electrostatic interactions, microdomain effects would be expected to be maximal within the membrane’s electric double layer that is approximately 1 nm at physiological ionic strength.

Therefore we sought to examine extent to which electrostatic interactions contribute to microdomains under steady state conditions. We found that the reaction rate coefficient had weak dependence on the enzymes’ distance from the pore surface, regardless of ionic strength, which strongly suggests microdomains arise from a different basis. Consistent with this argument is our recent finding that ionic-strength-dependent changes in the Ca^2+^ at the membrane surface has negligible impact on SERCA Ca^2+^ affinity [98].This confirms that localized substrate pools near the surface stem from non-equilibrium conditions, namely a net flux of substrate from the extracellular or cytosolic domains toward the membrane. For ATP, the steady-state flux toward the cytosolic side of the membrane could arise from the creatine and adenylate kinase shuttles [30, 93], while localized ATP gradients on the extracellular side could result from F10ATPase or translocase activity [8]. For ions such as Ca^2+^, membrane-localized gradients could arise from small inward fluxes of plasma membrane currents or leak from compartments such as the endoplasmic reticulum [97].

Our results suggest that the predominant effect of charging the membrane is to increase the concentration of ATP within the entire pore interior. This was evident in the predicted ATP concentration profile within the pore (see Fig. S1), which varied significantly from the bulk reservoirs. This raises an interesting possibility that NDA activity could be modulated through controlling the surface charge by varying membrane lipid composition. Such variations in lipid composition and surface charge are known to occur during phagocytosis [99] and within neural synapses [67].

In addition to substrate/surface interactions, we demonstrate that electrostatic interactions between nucleotide substrates and their enzymes targets accelerate NDA activity. Favorable long-range electrostatic interactions between enzymes and substrates are well known to optimize diffusion-limited reactions in biological systems [63]. Chiefly, enzymes that bear charges complementary to their substrates typically exhibit reaction rates that are several orders of magnitude higher than rates observed with neutral species or at high ionic strengths that shield electrostatic interactions [51]. We specifically address this for CD39, CD73 and charged membranes. CD39, for example, appears to have a slightly greater density of positively-charged amino acids near the nucleotide binding domain. We would expect this positive charge center to enhance the association rate via complementary electrostatic interactions through Arg56, Lys79, Lys80, and Lys82. This was determined by visual inspection of the electrostatic potential of a representative CD39 structure, the NTPDase2 from *Legionella pneumophila* (PDB code 4BR7 [24]) using the Adaptive Poisson Boltzmann Solver APBS[100]. Hence we expect this to facilitate the rapid reaction, though to our knowledge rates with respect to ionic strength for this enzyme have not been reported.

Beyond the role of electrostatics in shaping k_*on,ATP*_, our results demonstrate the kinetic advantage of co-localizing charged enzymes. When the enzymes were co-localized, the influence of electrostatic interactions on the reaction rates were most strongly evident, with the fastest rates reported for closely-opposed enzymes that adopt surface charges complementary to their substrates. This finding mirrors trends observed in other coupled enzymatic processes; namely in the event that enzymes or reactive sites are sequentially aligned for coupled enzymatic reactions, electrostatic channeling of substrates is commonly exploited in nature to optimize the rate or efficiency of substrate conversion [75, 101, 102]. As an example, a computational study of the DHFR-TS enzyme in prokaryotes has revealed that tetrahydrofolate production rates are accelerated by a patch of positively-charged amino acids between the thymidylate synthase and dihydrofolate reductase reactive sites, which facilitate transfer of the anionic dihydrofolate intermediate [21]. Significantly, when the enzymes’ charges were complementary to those of the reactants, the overall reaction efficiency exceeded predictions for the uncharged, confined enzymes and the enzymes in bulk solution. Hence, these variations should profoundly influence the dynamics and relative distribution of nucleotides within extracellular junctions.

### 4.4 Limitations and Future directions

In order to work with the system that was numerically solvable, we made several assumptions. Firstly, we assumed all enzymatic reactions were fast compared to the diffusion of nucleotides between reactive centers. NDAs are known to rapidly manage nucleotide pools with reaction rates on the order of 1 μM s^−1^ [31]. Since the intrinsic reaction rates of these enzymes vary quite considerably depending on the isoform and cell type, we assumed reaction-limited conditions for simplicity and generality. It may also be appropriate to consider feedback inhibition, given evidence that productions can hinder NDA-catalyzed AMP hydrolysis. We additionally assumed spherical shapes for the proteins; while this may seem to miss important details, our previous studies have indicated that the native structure of the proteins has a marginal influence on reaction dynamics [52]. We additionally assumed constant membrane potentials, although this can be expected to vary in real lipid systems such as during phagocytosis [99]. We additionally considered all reactions to be in steady state for ease of simulation and analysis for simplicity, though effects of micro domains expected to be most evident under non-linear, non-steady state conditions that permit significant ATP accumulation. We also limited our membrane potentials to modest ranges for which the linearized Poisson-Boltzmann equation was appropriate. For stronger potentials, the full PB equation would be more appropriate but also more computationally expensive [103]. Further, in highly charged and confined domains, the diffusing substrate can be expected to contribute to the shielding of electric charge, which advocates for the use of the Poisson-Nernst-Planck formalism (see [51]).

Although the enzyme and pore representations were simplistic in this study, our finite element modeling approach can be refined with detailed structural models of enzymes and their cellular environment. As an example, we have previously used finite element models to probe Ca^2+^ binding rates to myofilament proteins bound to actin chains [52, 53, 104] at atomic resolution, using mesh building software [105] applied to structures found in the Protein Databank. A similar approach could be used to probe the dynamics of other enzyme-catalyzed reactions in detailed molecular environments, such as those based on atomistic-resolution simulations of a crowded cytoplasm [106]. Along these lines, simulations of NDA activity in crowded synapses are warranted.

## 5 Conclusions

Sequentially-coupled enzymatic processes have been extensively probed in the literature. Our contribution in this paper complement these studies through offering insights into nucleotide distributions and NDA activity within extracellular junctions. Our results are consistent with the well-established notions of electrostatic channeling for accelerating reaction rates and for the benefits of co-localization. Our approach is unique in its basis in a finite-element framework that allows for the direct incorporation of electron or confocal microscopy data, such as for serial block images of neurons or neuromuscular junctions [107], or for atomistic resolution molecular structures of NDA s. In order to generalize our results, we utilized a simplified pore/spherical enzyme framework for which we could easily vary system parameters such as distances and radii, which could not be afforded with structurally detailed models. Based on our earlier studies, while details of the surface charge density of an active site can influence reaction rates, nuances of the protein structure generally do not substantially impact the results. This can be easily seen in our plots, which show relatively small changes in reactivity and efficiency as a function of significant changes in enzyme sizes or separation distances. This observation is helpful in reducing the spatial complexity of reaction simulations in complex media. As we demonstrated in [23], effective diffusion rates of small molecules in structurally-realistic crowded solutions did not significantly differ from those computed using perfect spheres of fixed radii at a similar packing fraction. This is in agreement with [52, 104, 108]. As shown in our results, this permits the contributions of crowders to diffusion-limited association rates to be accounted for in a rather simple implicit form, namely by accounting for configurational entropy of the reacting substrate.

These simulations of steady-state NDA nucleotide hydrolysis activity in nanoscale porous geometry mimic gap junctions between cells. We found that confinement and high charge densities within confined domains alter nucleotide concentrations relative to bulk, independent of NDA activity. Additionally, we demonstrate that for NDAs localized to the pore, confinement of NDA reduces activity relative to bulk, and the nature of surface interactions determines the advantage of co-localization on reaction efficiency. Of these, charges substantially influence reaction kinetics, particularly contributions of complementary membrane charge to ATP in the initial reaction.

We believe these findings provide new insights into the activity of purinergic receptors and other proteins that respond to extracellular ATP concentrations. Incorporating these features could expand our capacity to probe physiological phenomena *in vivo*, monitor and tailor drug delivery kinetics (reviewed in [109]), and engineer biosynthetic pathways, especially those utilizing immobilized enzymes [44, 91, 110]. Additional future directions include probing reaction dynamics relative to enzyme activity. We could further compare how the dynamic nucleotide signals influence the activity of purinergic receptors and ATPase activity on the extracellular domain, like F_1_-F_*O*_ ATP synthase[8].

## 6 Acknowledgements

Research reported in this publication was supported by the Maximizing Investigators’ Research Award (MIRA) (R35) from the National Institute of General Medical Sciences (NIGMS) of the National Institutes of Health (NIH) under grant number R35GM124977. We further acknowledge support from the American Chemical Society Petroleum Research Fund 58719-DN16. This work used the Extreme Science and Engineering Discovery Environment (XSEDE) [111], which is supported by National Science Foundation grant number ACI-1548562.

## 7 Method

### 7.1 Overview

The purpose of this study is to understand the steady-state properties of nucleotide hydrolysis by NDAs in nanoscale extracellular domains. This was achieved through computer simulations of electrokinetic transport, similar to those in [51], modified to handle reaction equilibria. The theoretical model includes partial differential equations defined on a continuum problem domain. These equations are solved numerically using the Finite Element Method. Steady-state conditions were assumed. The equations were solved for three-dimensional geometries resembling the porous materials in [51, 80]. Rather than attempting to simulate the intricate geometries of *in vivo* systems, the geometry of these porous materials was chosen in order to reduce the computational cost of the simulations, simplify interpretation of the results, and improve the reproducibility of our findings. The porous material is modeled as a thin membrane sepa-rating two aqueous reservoirs. A concentration gradient can be created between the two reservoirs to drive transport across the membrane. The simulations study diffusion taking place inside circular pores in the membrane. The computational model allows for the control of key geometric, electrostatic, and kinetic parameters, so that the effects of various phenomena can be resolved. Predicted substrate gradients that develop in the materials were used to estimate reaction rate coefficients, which directly relate to the kinetics of substrate transport. Additional physics was enabled in the computational model that accounted for electrostatic interactions and surface reactions. Chief outcomes of this study are 1) a computational model for molecular transport and 2) quantitative data for describing how confined domains and electrostatic interactions control molecular transport. For comparisons against bulk conditions, we assumed enzyme densities corresponding to millimolar concentrations.

### 7.2 Theory

In these simulations, the enzymes are held in fixed positions while the substrates are allowed to diffuse. The problem domain is approximated as a continuum, with the diffusing chemical species considered to be point particles. Under these circumstances, the reaction encounter distance is just the radius of the enzyme. The enzymes are therefore approximated as spheres.

Three species of substrate are included: A, AMP, and Ado. ATP is converted to the AMP product when it encounters the surface of CD39, followed by AMP’s conversion to adenosine on enzyme CD78:

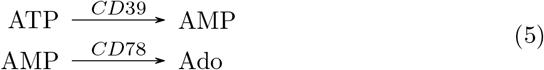

We define the concentration of a given species S as *c_S_*, which is an unknown spatial function to be found by solving the governing diffusion equation. The ion flux of species S is a vector field, 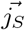, related to the change in concentration with respect to time through a continuity equation,

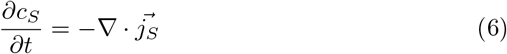

The diffusion of ions in a fixed electrostatic field is described by the Smolu-chowski equation [112], where the flux includes both a Fickian diffusion term and a term due to the electrostatic force:

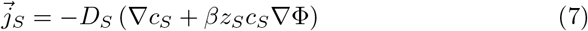

Where *z_S_* is the electric charge of species S, Φ is the electric potential as a scalar field, *D_S_* is the Fickian diffusion coefficient for species S in the relevant media, and *β* is 1/*k_B_T* for temperature *T* and Boltzmann constant *k_B_*. In this equation, the diffusion coefficient is assumed to be homogeneous and isotropic.

Using this flux in the continuity equation, the Smoluchowski equation can be written as:

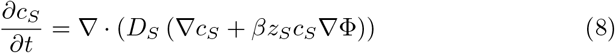

Under steady state conditions the concentration of species S does not vary in time, and so the governing differential equation is:

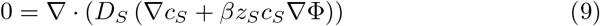

To reduce the computational burden, an alternate form of the Smoluchowski equation is used. The substrate flux is expressed as:

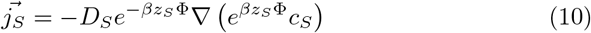

The equivalence of these two expressions for the flux can be readily verified using the product rule for gradients. The advantage of this alternate form is that it allows for the application of the Slotboom transformation [113] [114]:

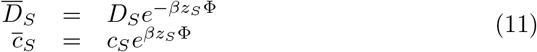

After applying this transformation, the Smoluchowski equation is expressed in a form analogous to a simple Fickian diffusion equation:

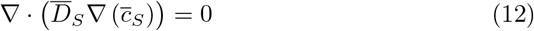

Equation 12 must be solved for each species.

We also define the integrated flux over any surface Γ as

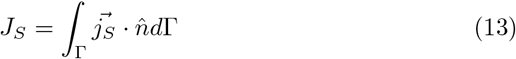

where 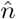 is the unit normal to the surface Γ.

The reaction kinetics at an enzyme are assumed to follow a simple rate law. For the reactions in Equation 5, the rate laws are given by

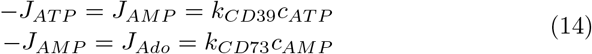

which defines reaction rate coefficients *k*_*CD*39_ = k_*on,ATP*_ = k_*prod,AMP*_ and *k*_*CD*73_ = k_*on,AMP*_ = k_*prod,Ado*_.

The calculation procedure begins by solving Equation 12 for 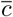 for each species, then computing the flux vector fields from the Slotboom transformation of Equation 7, then integrating the flux over the enzyme surface, and using the integrated flux to calculate the rate coefficient. The rate laws of Equation 14 are enforced at each enzyme by requiring that 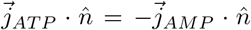 on the surface of CD39, and 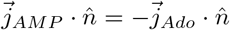 on the surface of CD73.

A summary of concentration boundary conditions applied to the model is presented in Table 1.

The solution of Equation 12 requires knowledge of the electric potential Φ throughout the model. The electric potential is found by solving the linearized Poisson-Boltzmann equation:

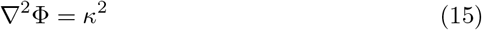

where *κ* is ionic strength which is the inverse of the Debye length. For pure aqueous solvent, *κ* = 0 and therefore Eq. 15 reduces to the Poisson equation commonly used in electrostatics.

### 7.3 Numerical approach

Methodologies are generally as described in Sun et al., [51].

The system of partial differential equations and boundary conditions described above was solved numerically using the Finite Element Method. The open-source finite element package FEniCS[115], version 2017.2.0 was used to conduct the simulations. This software is publicly available at fenicsproject.org.

A second-order polynomial (Lagrange) basis set was used for all finite elements. The differential equations to be solved were all linear, so no nonlinear solution schemes were required. Various linear solvers and preconditioners were employed in order to obtain solutions.

Python-based analysis routines were used to set up, solve, and post-process the finite element models. All code written in support of this publication is publicly available at https://bitbucket.org/pkhlab/pkh-lab-analyses. Simulation input files and generated data are available upon request.

## Supplementary Information

### S.1 Tables

**Table S1:** Summary of cases run

| Fig (Sect) | qA/EA | qB/Eb | Pore | Crowders | $\lambda_D$ |
| --- | --- | --- | --- | --- | --- |
| Fig. ?? (1) | -1/0 | +1/0 | 0 | 0 |  |
| 2 | -1/+1 | +1/-1 | var. | 0 | ? |
| 3 | -1/0 | +1/0 | 0 | var. | ? |
| Fig. ??c | -1/0 | var./0 | 0 | var. |  |

### S.2 Figures

**Figure S1:**
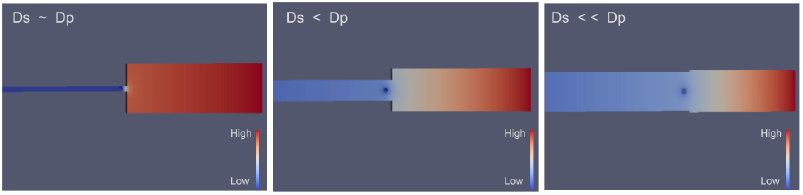
Concentration profile for three different values of pore radius.

**Figure S2:**
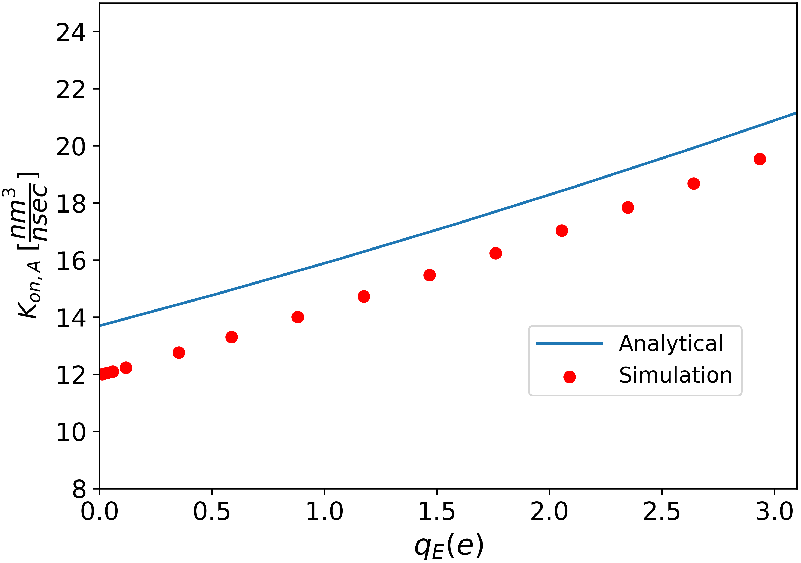
A comparison between theoretical (see Eq. 2) and simulation result which validates our model of simulation.

**Figure S3:**
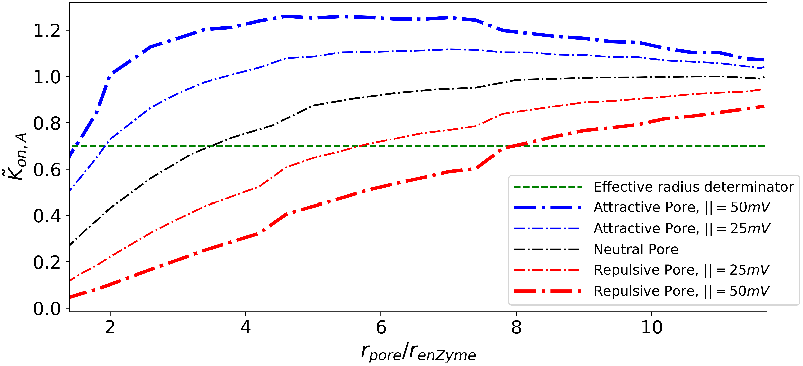
Effective pore radius for larger relative size of the pore to enzyme radius.

**Figure S4:**
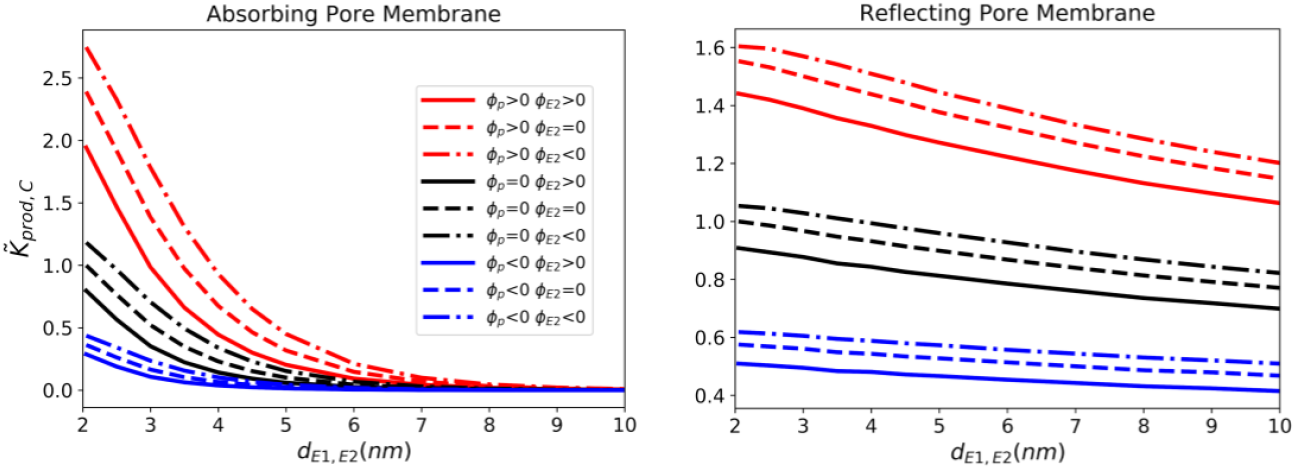
Effects of charge. Effect of charge sign composition on reactivity, given *z_ATP_*=−1, *z_AMP_*=+1, *z_Ado_*=0 and Φ_*CD*39_ > 0 Normalized reaction rate coefficient for production of Ado as a function of distance between enzymes.

**Figure S5:**
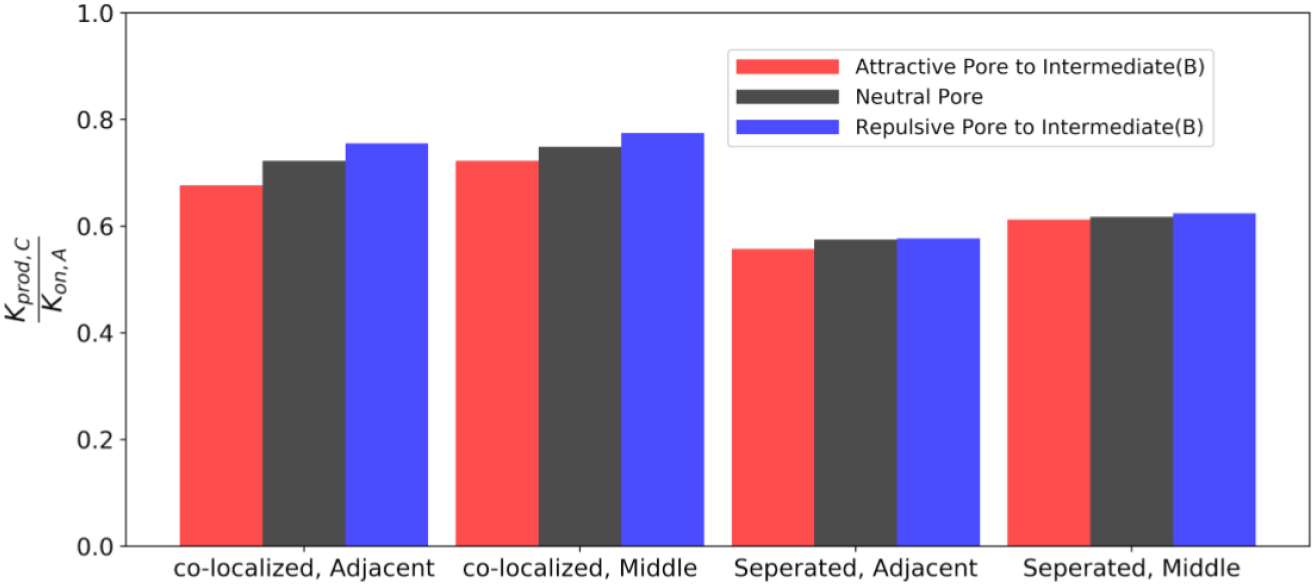
Effects of charge. Efficiency of sequential enzymes for different pore wall electric potentials, given *z_ATP_*=−1, *z_AMP_*=+1, *z_Ado_*=0 and Φ_*CD*39_ > 0

**Figure S6:**
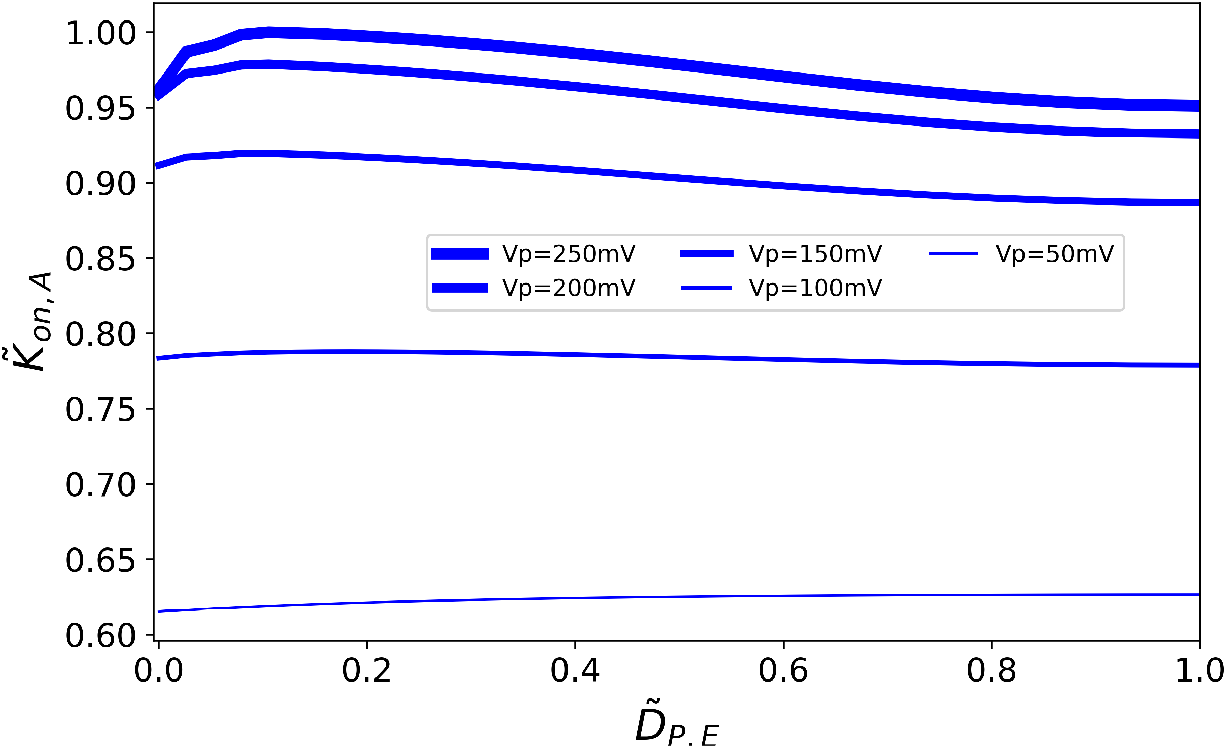
For large potential of the pore in comparison with enzyme potential, as the enzyme get close to the pore membrane, due to the attraction between pore membrane and substrate ATP the concentration of ATP is more and so it leads to an increase to the reactivity of enzyme. However, when the enzyme is getting too closer to the pore membrane, the competition between pore membrane and enzyme, especially for very large potential on membrane, is increasing, leading to a sudden decrease in reactivity.This obtained for kappa=1, and Dp=5DE.

**Figure S7:**
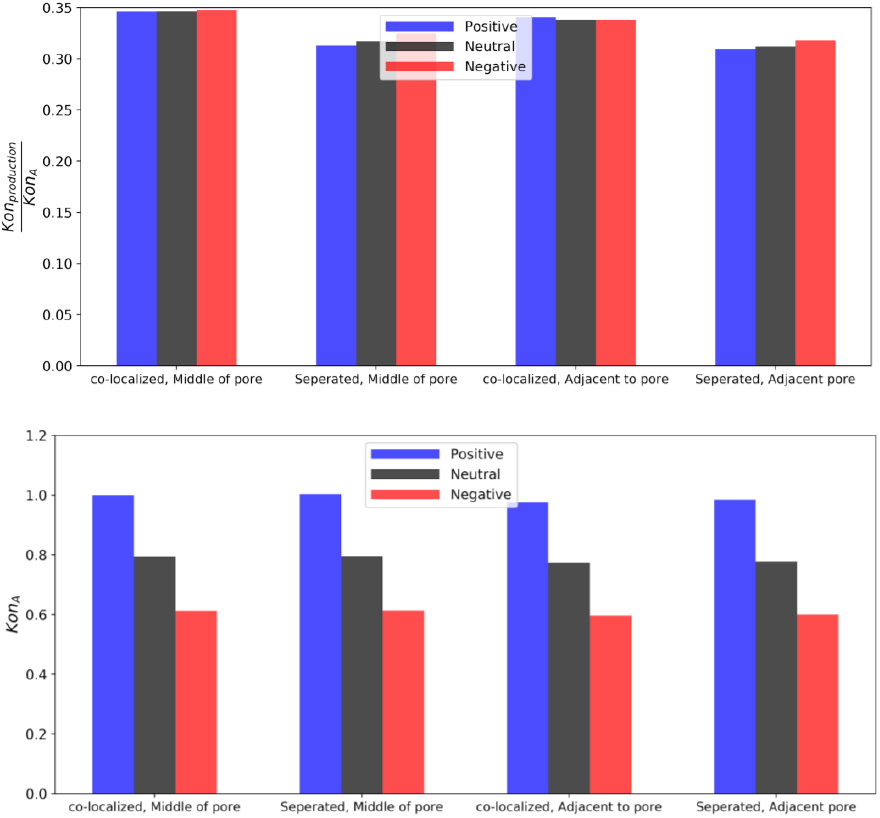
Coloc/tethering effects on keff. These data obtained for the case which maximize the efficiency with *VE*1 > 0, *VE*2 < 0, Dp=8 and Vp=25mV. The konA does not change when enzymes get close to the wall for all different charges of the pore wall. However, I have tried a case when Dp=11, kappa=1 and Vp=100 and saw that the konA increase as the enzymes get close tho the wall for attractive case between species ATP and pore wall (here positive), although the difference is not considerable(around 1.5 percent change). There is two factors affecting konA when it get close to the wall: The capacitance change when the enzyme get close to the wall. However, the concentration of ATP species is more near the wall of the pore.

**Figure S8:**
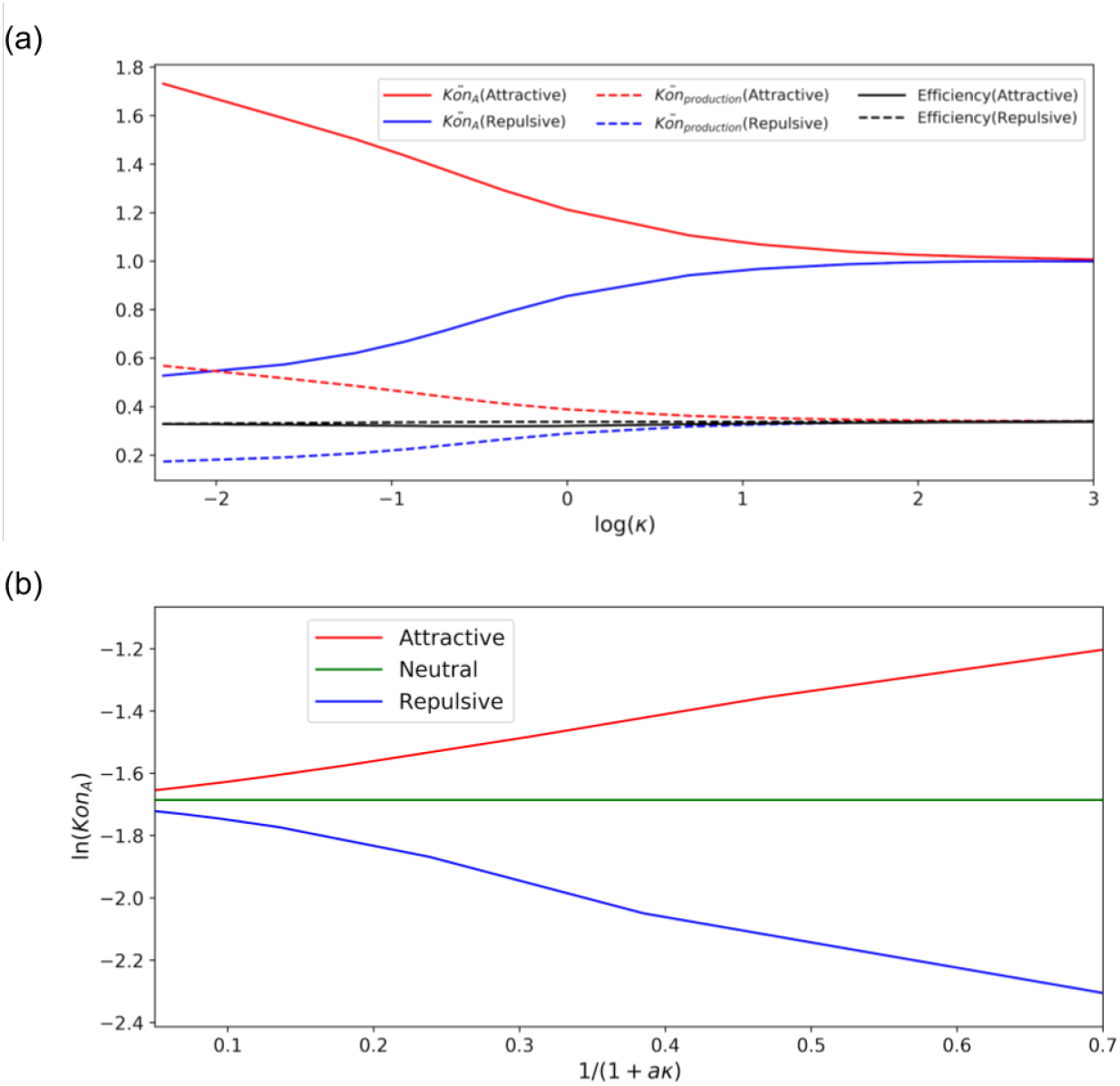
Effect of Debye Length on reactivity of sequential enzyme: a) Reactivity of the first enzyme and second enzyme as a function of pore to enzyme distance for electrostatical compositions with maximum (red) and minimum (blue) reactivity along with the efficiency (black) c) b)effective pore radius based on potential of the pore and Debye length. The green curve shows the data for uncharged reactivity of the first enzyme as a function of real physical size. based on the data we obtained from charged simulation, we can match an effective pore size which for attractive is bigger and for repulsive is less than its real size.

